# Bovine blastocyst development depends on threonine catabolism

**DOI:** 10.1101/397562

**Authors:** V Najafzadeh, H Henderson, R Martinus, B Oback

## Abstract

Increasing evidence suggests that pluripotency is a metabolically specialised state. In mouse, inner cell mass (ICM) cells and ICM-derived pluripotent stem cells (PSCs) critically depend on catabolising the amino acid threonine, while human PSCs require leucine, lysine, methionine or tryptophan. However, little is known about the specific amino acid requirements of putative pluripotent cells in bovine. We selectively depleted candidate essential amino acids (EAAs) from individually cultured bovine embryos to study their role in blastocyst development. Depleting one (-T, -M), two (-MT, -CM, -CT, -IL, -IK, -KL) or three (-CMT, -IKL) EAAs from chemically defined protein-free culture medium did not affect the morula-to-blastocyst transition from day five (D5) to D8 *in vitro*. By contrast, removing six (-CIKLMT, -FHRYVW), nine (+CMT, +IKL), eleven EAAs (+T, +M) or all twelve EAAs increasingly impaired blastocyst development. As no clear candidate emerged from this targeted screen, we focussed on threonine dehydrogenase (TDH), which catalyses threonine catabolism. *TDH* mRNA and protein was present at similar levels in trophectoderm (TE) and ICM but absent from several adult somatic tissues. We then treated morulae with an inhibitor (Qc1) that blocks TDH from catabolising threonine. Continuous exposure to Qc1 reduced total and high-quality blastocyst development from 37% to 26% and 18% to 8%, respectively (P<0.005). This was accompanied by ∼2-fold decrease in ICM, TE and total cell numbers (P<0.005), which was due to increased autophagy (P<0.05). At the same time, ICM-(*NANOG*) and TE-restricted (*KRT8*) genes were up-and down-regulated, respectively (P<0.05). In summary, bovine blastocyst viability depended on TDH-mediated threonine catabolism. However, ICM and TE cells did not metabolically differ in this regard, highlighting species-specific connections between metabolism and pluripotency regulation in mouse vs cattle.

## INTRODUCTION

Following fertilization, the diploid mammalian zygote undergoes a series of DNA replications and cleavage divisions, partitioning its cytoplasm into progressively smaller nucleated daughter cells, called blastomeres, without increasing total embryo volume. Energy production during these cleavage divisions is largely aerobic with pyruvate as the preferred substrate to generate ATP from oxidative phosphorylation [1]. During *in vitro* culture, early preimplantation embryos have remarkably simple nutritional needs, requiring only physiological salt solutions, an energy source and a fixed nitrogen source. In chemically defined media, these include a few key ingredients, such as pyruvate, lactate, endogenous fatty acids and amino acids [1].

After a few cleavage divisions, the blastomeres compact and form a tight morula. This structure is unique to placental mammals and comprises an outer layer and an internalized cell mass. The outer cells form an epithelium, channelling water and ions inside to form a small fluid-filled cavity, the blastocoel. Cavitation marks the appearance of an early blastocyst, a hollow ball of cells with an outside trophoblast layer, often called trophectoderm (TE), enclosing a small inner cell mass (ICM) to one side of the expanding blastocoel. The trophoblast gives rise to fetal placenta, while the ICM segregates into the hypoblast, which form extraembryonic yolk sac endoderm, and the pluripotent epiblast, which develops into all intraembryonic lineages, including germ cells, and extraembryonic yolk sac mesoderm. In mouse and rat, isolated epiblast is also the source of naïve pluripotent stem cells (PSCs) that can be permanently cultured *in vitro* [2].

During these developmental processes, glucose becomes an important substrate and oxygen consumption rises, accompanied by a sharp increase in glycolysis. These metabolic changes are largely due to the energy demands of the Na+, K+ ATPase pump that is required for blastocoel formation and of increasing protein synthesis [3]. They coincide with the end of cleavage divisions, initiation of net cell growth, replicative proliferation and stable nucleo-cytoplasmic ratios [4].

In the absence of other protein sources, the precise combination and concentration of amino acids that is required for cultured bovine embryos at different developmental stages is still a matter of debate [5, 6]. However, there is little evidence that individual amino acids could be harmful, especially at near-physiological concentrations. Therefore, optimised culture media usually include the full complement of all 20 standard amino acids, ideally in a composition closely resembling the reproductive tract fluids in which embryos normally develop [7]. During *in vitro* development, this composition may have to change according to changing embryo needs. For example, individual amino acids are depleted at higher rates at later developmental stages, implying increased demand as development progresses [8]. Profiling such amino acid turnover has been used to predict developmental competence of embryos from a variety of species, e.g., human, pig and cattle (reviewed in [9]).

Conventionally, amino acids have been categorised into ‘essential’ and ‘non-essential’ depending on whether they can be synthesized by the whole organism or not, respectively. Regardless of this distinction, all 20 standard amino acids are required for protein synthesis and are usually supplied in the culture medium of early embryos [6]. This supply is either direct, as free amino acids, or indirect, as serum or purified serum albumin that can be internalized by embryos and degraded intracellularly [10]. Once internalized, amino acids and their complex interactions play a number of overlapping metabolic and cellular roles. Quantitatively the most important one is protein synthesis, which is directly measured through incorporation of exogenous radiolabelled amino acids into protein of preimplantation embryos [11]. Amino acids are also involved in numerous other biosynthetic pathways, including nucleotide synthesis, oxidation in the Krebs cycle, protection against oxidative stress, osmotic and pH regulation, and one-carbon metabolism (reviewed in [7]).

One essential amino acid with exceptional metabolic flexibility is threonine. Apart from protein translation and post-translational modification, such as phosphorylation and O-linked glycosylation, threonine can be enzymatically catabolized to produce energy, lipids, nucleotides and one-carbon donors [12]. This wide range of roles depends largely on the activity of threonine dehydrogenase (TDH), a mitochondrial enzyme that catalyses the two-step breakdown of threonine into acetyl-CoA and glycine, feeding the Krebs cycle and one-carbon metabolism, respectively. Using systematic amino acid depletion, threonine was identified as being critically important for epiblast-derived mouse PSCs [13]. In the absence of threonine, thymidine biosynthesis and cell cycle progression were impaired while differentiation was accelerated [13, 14]. Furthermore, metabolism of threonine and S-adenosylmethionine (SAM), the main methyl-donating molecule, were closely coupled in PSCs [15]. Decreased threonine levels led to a concomitant decrease in SAM levels, which manifested as a decrease in specific histone trimethylation levels [15]. Inhibition of TDH activity by quinazolinecarboxamide (Qc) compounds specifically impeded cell growth and induced autophagy of mouse PSCs [16]. These pharmacological studies were supported by genetic TDH-overexpression and -knockdown experiments that resulted in enhanced and reduced reprogramming efficiency, respectively, into induced PSCs (iPSCs) [17]. Collectively, these findings demonstrated the important role of TDH-mediated threonine catabolism in regulating pluripotency. In humans, TDH is predicted to be an inactive pseudogene, however, this is based on three different mutations present in a small sample size and no functional TDH assay [18]. Human PSCs rely on the uptake of extracellular methionine for SAM production as threonine cannot be used for SAM production [13], which instead use methionine [19]. Consequently, methionine deprivation leads to loss of histone methylation and apoptosis [19]. A similar effect was observed after knockdown of methionine adenosyltransferases that catalyse the conversion of methionine into SAM, suggesting that maintenance of SAM levels, rather than methionine, was critical for cell survival. Methionine deprivation also led to a rapid and specific decrease in histone trimethylation [19]. Taken together, threonine and methionine are critical for PSC survival through modulating SAM concentrations, potentially linking the metabolic and epigenetic pathways in PSCs.

Here we investigated the role of threonine and methionine in bovine blastocyst development with a particular focus on forming the ICM, the founder tissue for embryo-derived putative PSCs. We show that blocking TDH activity leads to increased autophagy in bovine blastocysts, demonstrating a requirement of threonine catabolism for embryonic cell viability in cattle.

## MATERIAL AND METHODS

Chemicals were purchased from Sigma-Aldrich (Auckland, New Zealand) unless indicated otherwise.

### Animal Studies

All animal studies were undertaken in compliance with New Zealand laws and were approved by the Ruakura Animal Ethics Committee.

### Cell lines and tissues

Mouse embryonic fibroblasts (MEFs) were derived from day 13.5 embryos (Crl:CD/ICR strain) as described [20] and cultured in DMEM/F12 with 10% fetal bovine serum (FBS) and 1x MEM non-essential amino acids. Muscle fibroblasts were isolated from the quadriceps femoris of adult CD mice and cultured under the same conditions as MEFs. Male v6.5 ESCs (C57BL/6 × 129/SV strain) were cultured feeder-free on 0.1% gelatine in DMEM/F12 with 20% FBS, 100 µM β-mercaptoethanol, 1x MEM non-essential amino acids, 2000 U/ml of recombinant human LIF (20 ng/ml) and 0.4 µM PD0325901. Following doxycycline (Dox) - dependent induction, secondary iPSCs were derived from homozygous R26^*rtTA*^, Col1a1^4F2a^ MEFs (Jackson laboratory, stock no. 011004) and maintained in ESC culture medium as described [21].

### TDH-overexpressing cells

For stable *TDH* overexpression, rejuvenated bovine embryonic fibroblasts (BEFs), constitutively expressing the Tet-On 3G reverse tetracycline transactivator (manuscript in preparation), were transfected with an inducible *Piggybac* (*pB*) response vector for bovine *TDH* expression (manuscript in preparation). Controlled by a Dox-inducible P_TRE3G_ promoter, the full-length *TDH* open reading frame was fused to a C-terminal *MYC*-tag and co-expressed a *puromycin* selection marker, via an internal ribosome entry site (insert synthesized by GENEART, Germany). Stable BEF5-TDH cells were generated by using Lipofectamine^®^LTX/PLUS^™^, (ThermoFisher Scientific, New Zealand) to co-transfect BEF5 cells with *pB_TRE_TDH_MYC_PuroR* and the *pCyL43* transposase [22] according to the manufacturer’s instructions. Following induction with Dox (2 µg/ml) for 48 hours, cells were puromycin-selected (2 µg/ml) in the continuous presence of Dox for several days. Stable populations of puromycin-resistant cells were cryopreserved and characterised for transgene expression.

### In vitro Production (IVP) of Bovine Embryos

*In vitro*-matured (IVM) oocytes from a batch of slaughterhouse ovaries of mixed breed dairy cows were fertilized with frozen–thawed semen from a sire with proven *in vitro* fertility as described [23]. For cumulus-free cultures, the corona was dispersed after IVF for 22–24 h by vortexing oocytes in 500 µl of 1 mg/ml bovine testicular hyaluronidase in Hepes-buffered SOF (HSOF), followed by two washes in HSOF. For *in vitro* culture (IVC), 10 embryos were pooled in 20 µl of early SOF (ESOF) medium and cultured for five days (D0: fertilization). On D5, embryos were categorised into three groups: i) 1-cells, ii) less than 8-cells and iii) equal/more than 8-cells. Embryos from each group were then randomly distributed across the desired number of treatments by using ‘proportionate’ pooling, whereby each group supplied equal numbers of embryos per treatment but the number per treatment varied among groups. Care was taken that the number of embryos allocated for each treatment was approximately equal. Embryos were washed in 2 ml of pre-warmed PBS with 0.1 mg/ml cold soluble PVA (M_r_: 10-30000) (‘PBS/PVA’) and 20 µl drops of late SOF (LSOF) to remove traces of amino acid/protein carry-over before transfer into 5 µl LSOF drops containing 10 μM 2,4-dinitrophenol [24] for single embryo culture. For dropout experiments, EAAs, all NEAAs, BSA (8 mg/ml) and Glutamax (1 mM) were first omitted from LSOF (basal LSOF or bSOF), before reintroducing individual components according to the different treatment groups. Biological replicates were run on different experimental days, using different batches of ovaries and embryos.

| **Amino acid (abbreviation)** | **Category** | **Concentration (mM)** |
| --- | --- | --- |
| L-Arginine-HCl (R)* | semi-essential | 0.60 |
| L-Cystine (C)* | semi-essential | 0.10 |
| L-Histidine (H) | essential | 0.20 |
| L-Isoleucine (I) | essential | 0.40 |
| L-Leucine (L) | essential | 0.40 |
| L-Lysine monohydrochloride | essential | 0.40 |
| L-Methionine (M) | essential | 0.10 |
| L-Phenylalanine (F) | essential | 0.20 |
| L-Threonine (T) | essential | 0.40 |
| L-Tryptophan (W) | essential | 0.05 |
| L-Tyrosine (Y)* | semi-essential | 0.20 |
| L-Valine (V) | essential | 0.40 |

\*These amino acids are semi-essential as arginine is often synthesized in inadequate amounts in the urea cycle, while cystine, the oxidation product of cysteine, and tyrosine can be synthesized from methionine and phenylalanine, respectively.

For inhibitor experiments, LSOF was supplemented with TDH inhibitor Qc1 (5 mM stock, 50 μM final concentration) or equimolar dimethyl sulfoxide (DMSO) solvent controls. Embryo cultures were overlaid with mineral oil and kept in a humidified modular incubation chamber (ICN Biomedicals Inc., Aurora, OH) gassed with 5% CO_2_, 7% O_2_, and 88% N^2^. Embryos were morphologically graded on D7 or D8 as described [25].

### IVP of Mouse Embryos

Female mice (B6C3 or Swiss strain) at 8-12 weeks of age were intraperitoneally injected with 10 international units (IU) of pregnant mare serum on day zero at 5:00 pm. Human chorionic gonadotropin (10 IU/ml) was injected 48 hours later. After 14-17 h, mice were euthanized and oocytes recovered from the ampulla of each ovary according to standard procedures. Following washes in Hepes-buffered CZB medium with 0.1% PVP, oocytes were placed into pre-warmed bovine testicular hyaluronidase (1 mg/ml) for 15 min at 37°C to dislodge cumulus cells.

Subsequently, they were washed and cultured in M16 medium 20 µl culture drops and incubated at 5% CO_2_ and 37°C. After 5 h, embryos were artificially activated in calcium-free M16 with 4 mg/ml BSA (Cat.# 81-003-03), vitamins (Cat.# R7256), 5 µg/ml cytochalasin B and 1 mM SrCl_2_ After 6 h, cells were washed out in a new plate of M16 and spread out at 15-20 embryos per 20 µl drop. The next morning, parthenogenetically activated embryos were washed in pre-warmed PBS-PVA and randomly allocated to their respective treatments. For dropout experiments, EAAs, NEAAs and BSA were omitted from home-made M16 (basal M16 or bM16), before reintroducing individual components according to the different treatment groups. For inhibitor experiments, M16 was supplemented with 50 μM Qc1 or equimolar DMSO solvent controls. Embryos were individually cultured in 5 µl culture drops and morphologically graded on D4.

## Differential Staining

Blastocysts were treated with pronase to remove the zona pellucida [26] and exposed to a 1:4 dilution in HSOF of rabbit anti-bovine whole serum (B3759) for 45 min, rinsed in HSOF with 0.1 mg/ml PVA (HSOF-PVA), and placed into a 1:4 dilution in HSOF of guinea pig complement (S1639) containing 5 μg/ml propidium iodide and 40 μg/ml Hoechst 33342 for 15 min. After rinsing in HSOF-PVA, the embryos were mounted (ProLong™ Diamond antifade mountant; ThermoFisher Scientific) on glass slides and examined using an epifluorescence microscope (AX-70; Olympus). Blue and pink colours were designated as ICM and TE cells, respectively [27], and cell numbers were quantified the same day using ImageJ software.

### Autophagy

A commercial autophagy detection kit (Abcam, ab139484) was adapted for bovine embryos. Briefly, blastocysts were washed once with assay buffer, stained for 30 min at 38.5°C in the dark, washed again with 100 μl assay buffer and mounted in 5 μl of ProLong^®^ Diamond antifade. Stained embryos were analysed by epifluorescence microscopy.

### ICM and TE Isolation

For ICM isolation, blastocysts were treated with pronase to remove the zona pellucida [26]. They were then incubated in 0.2% TX-100 (w/v) for about 3 seconds and then transferred to PBS/PVA (0.1 mg/ml) buffer. By gently pipetting up and down, TE cells started dislodging while the ICM remained intact. For mechanical TE isolation, zona-free blastocysts were washed in PBS-PVA and protein-free HSOF-PVA before attaching them to the lid of a Petri dish in protein-free HSOF. A microsurgical splitting blade (ESE020; Bioniche Animal Health Inc.), mounted to a three-axis oil hydraulic hanging joystick micromanipulator (MO-188; Nikon Narishige), was lowered to separate the ICM from the majority of the TE tissue. Separated tissues were washed in HSOF and processed for further experiments.

### RNA Extraction and RT-PCR

Pools of grade 1-2 blastocysts were lysed in 10 µL *RNA*GEM™ Tissue *PLUS* (containing 0.5 µL *RNA*GEM™) and cDNA synthesized as described [28]. Reverse transcriptase was omitted in one sample each time a batch was processed for cDNA synthesis. For quantitative PCR (qPCR), a LightCycler (Roche) was used with primers shown in Supplemental Table 1. Primers were designed using NCBI/ Primer-BLAST and synthesized by Integrated DNA Technologies (IDT, IA, USA). All the reactions were performed either with the LightCycler FastStart DNA MasterPLUS SYBR Green I kit (Roche) or Takara Bio SYBR Green Master Mix (Clontech). The ready-to-use Hot Start LightCycler reaction mix consisted of 0.4 μl of each primer (10 μM), 2.0 μl LightCycler SYBR Green I master mix or Takara Master Mix, 6.2 μl diethylpyrocarbonate-treated water, and 1 μl cDNA template. The following four-segment program was used: 1) denaturation (10 min at 95°C), 2) amplification and quantification (20 sec at 95°C, 20 sec at 55°C–60°C, followed by 20 sec at 72°C with a single fluorescent measurement repeated 45 times), 3) melting curve (95°C, then cooling to 65°C for 20 sec, followed by heating at 0.2°C/sec to 95°C while continuously measuring the fluorescence), and 4) cooling to 4°C. Product identity was confirmed by gel electrophoresis and melting curve analysis. For relative quantification, external standard curves were generated from serial 10-log dilutions for each gene in duplicate or triplicate. One high-efficiency curve (3.6 ≥ slope ≥ 3.1, R^2^> 0.99) was saved for each target gene and imported for relative quantification compared to 18S RNA or GAPDH as described [28].

**TABLE 1.**
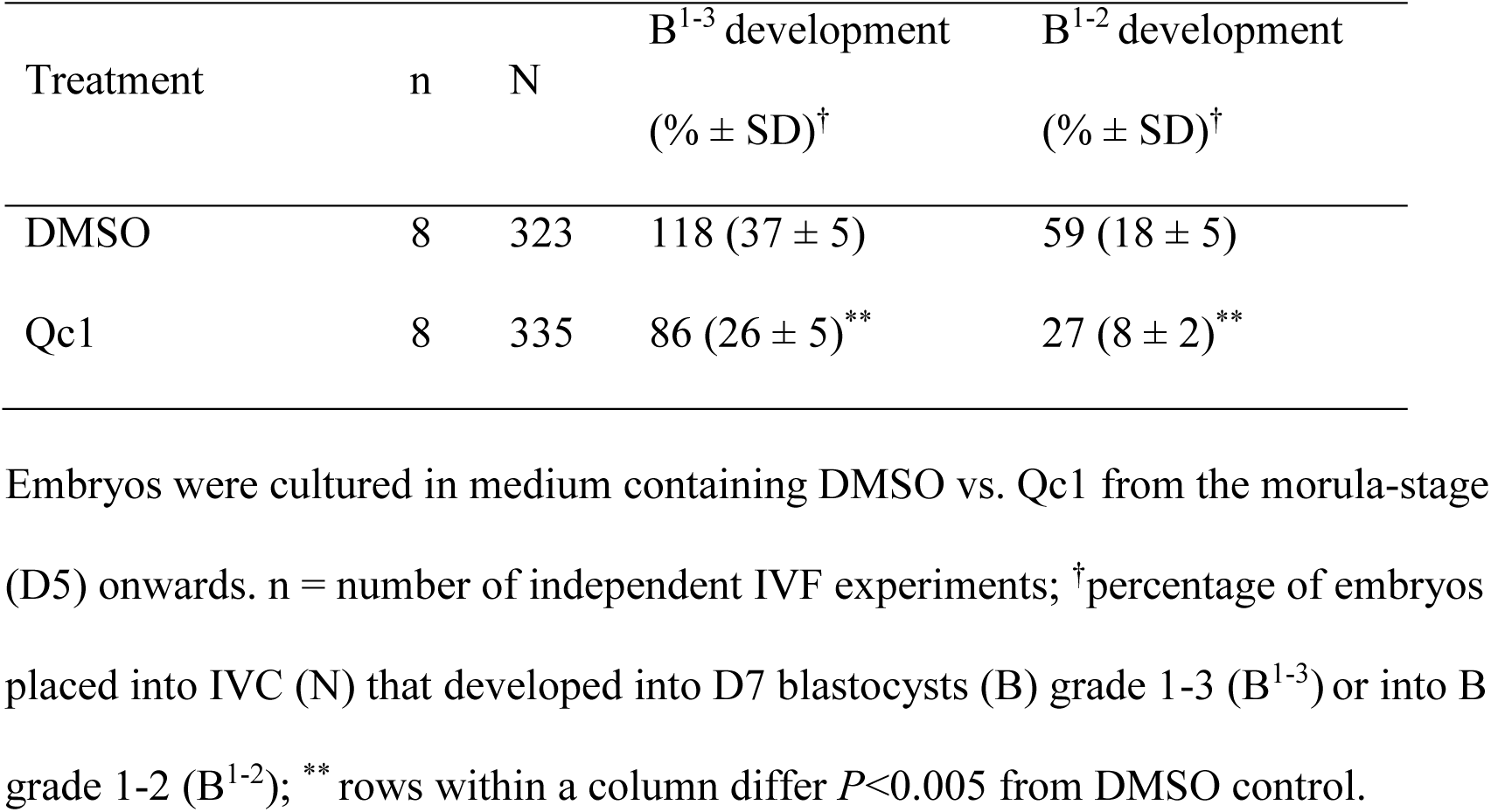
Effect of TDH inhibition on bovine blastocyst development. Embryos were cultured in medium containing DMSO vs. Qc1 from the morula-stage (D5) onwards. n = number of independent IVF experiments; ^†^percentage of embryos placed into IVC (N) that developed into D7 blastocysts (B) grade 1-3 (B^1-3^) or into B grade 1-2 (B^1-2^); ^**^ rows within a column differ *P*<0.005 from DMSO control.

### Immunofluorescence

Cells were fixed in 4% paraformaldehyde (PFA) for 15 min at 4°C, washed in PBS, quenched in 50 mM NH4Cl in PBS for 10 min, permeabilized in 0.1% (v/v) Triton X-100 in PBS for 10 min at room temperature and blocked in 5% donkey serum, 5% BSA in PBS for 30 min. Primary TDH antibody was used at 1:500 dilution, incubated overnight at 4°C, washed in PBS and detected with Alexa Fluor^®^ 488 or 546 donkey anti-rabbit secondary IgG antibodies (ThermoFisher Scientific) for 30 min at 38.5°C. All antibodies were diluted in blocking buffer. DNA was counterstained with 5 µg/ml Hoechst 33342. Preparations were washed in PBS before mounting (ProLong® Diamond antifade mounting medium). Negative controls were processed the same way, except that the primary antibodies were replaced with blocking buffer. Images were taken on an epifluorescence (Olympus BX50) and captured with a Spot RT-KE slider camera (Diagnostics Instruments Inc., MI, USA) using Spot software (version 4.6) or on a confocal microscope (Olympus FluoView FV1000).

### Mitochondria staining

Embryos were washed three times for 5 min each in 3% pre-warmed BSA-PBS. They were then incubated for 30 min in the same buffer containing 250 nM Mitotracker Red CMXRos (Molecular Probes M-7512, Oregon, USA) under 5% CO_2_ at 38.5°C. Subsequently, embryos were washed three times in pre-warmed PBS and fixed with 4% PFA for 30 min at RT before processing for TDH immunofluorescence as above.

### Quantification of Immunofluorescence

Epifluorescence images were acquired with all microscope and capture settings kept constant between technical replicates. Pixel intensity of single confocal frames from randomly selected nuclei was quantified (Olympus FluoView FV1000 with FV10-ASW 1.4 software). First, one random cytoplasmic area was background-subtracted from each image. The nucleus was marked as the region of interest, and series analysis was performed to compute the area and average intensity of the entire stack. Within each stack, the frame of highest average pixel intensity for the antibody channel was selected and divided by the corresponding Hoechst 33342 pixel intensity.

### Western Blotting

Western blot analyses were carried out using TDH and H3 (Abcam H3, ab1791) antibodies, both diluted 1:5000. Whole cell lysates (40 µg per lane) were resolved on 4–12% Bis-Tris SDS-Page gradient NuPage gels, transferred to nitrocellulose membranes and wet-blotted. The secondary antibody, goat anti-rabbit IgG-HRP (Dako, P0448), was used at 1:5000 and peroxidase activity was visualized with Western Lightning Plus-ECL kit (PerkinElmer, NEL105001EA).

### Statistical Analysis

Statistical significance was accepted at P<0.05. For blastocyst development, significance was only accepted when two different methods both gave significant results, namely pairwise Fisher’s exact test comparisons and Residual Maximum Likelihood analysis in Genstat 18 by culture condition, specifying run as a random effect. Student 2-tailed t-tests were used for average cell numbers and gene expression measurements. All bars represent standard error of mean (SEM). Unless stated otherwise, “N” denotes the number of samples analysed; “n” denotes the number of replicate experiments.

## RESULTS

### Bovine blastocyst development did not critically depend on candidate amino acids

We first established culture conditions for investigating the dependence of ICM development on individual amino acids. As BSA provides a major source of various amino acids for developing embryos, it was omitted from the culture medium from the 32-64 cell stage (D5) onwards. Absence of BSA reduced total blastocyst development under group culture conditions, and omission of NEAAs or EAAs further exacerbated this effect, in particular for high-grade blastocysts (Fig. S1A). As embryos themselves may be a confounding source of amino acids during group culture, we studied the effect of BSA and NEAA vs EAA dropout during single embryo culture. Under these conditions, there was a more severe reduction in blastocyst development when both NEAAs and EAAs were removed in the absence of BSA (Fig. S1B). To further refine the system, embryos were thoroughly washed in PBS/PVA before transfer into the dropout medium and single culture.

This further reduced development in the absence of BSA, NEAAs and/or EAAs (Fig. S1C). In the absence of any exogenous protein source for three days, only 6% of embryos developed into poor quality blastocysts on D8. Since dropping out EAAs had a more detrimental effect on bovine blastocyst development than dropping out NEAAs, we decided to deprive individually cultured embryos of candidate EAAs from D5 to D8 in subsequent experiments.

Mouse embryos and ESCs critically rely on threonine catabolism for survival. We therefore tested threonine dependency of bovine blastocysts using our previously established minimal culture conditions. Blastocyst development was not significantly affected by the absence of threonine compared to full medium, even when all NEAAs were dropped out at the same time (Fig. 1A). Reciprocally, threonine provision did not rescue blastocyst development when all other EAAs were dropped out.

**Figure 1.**
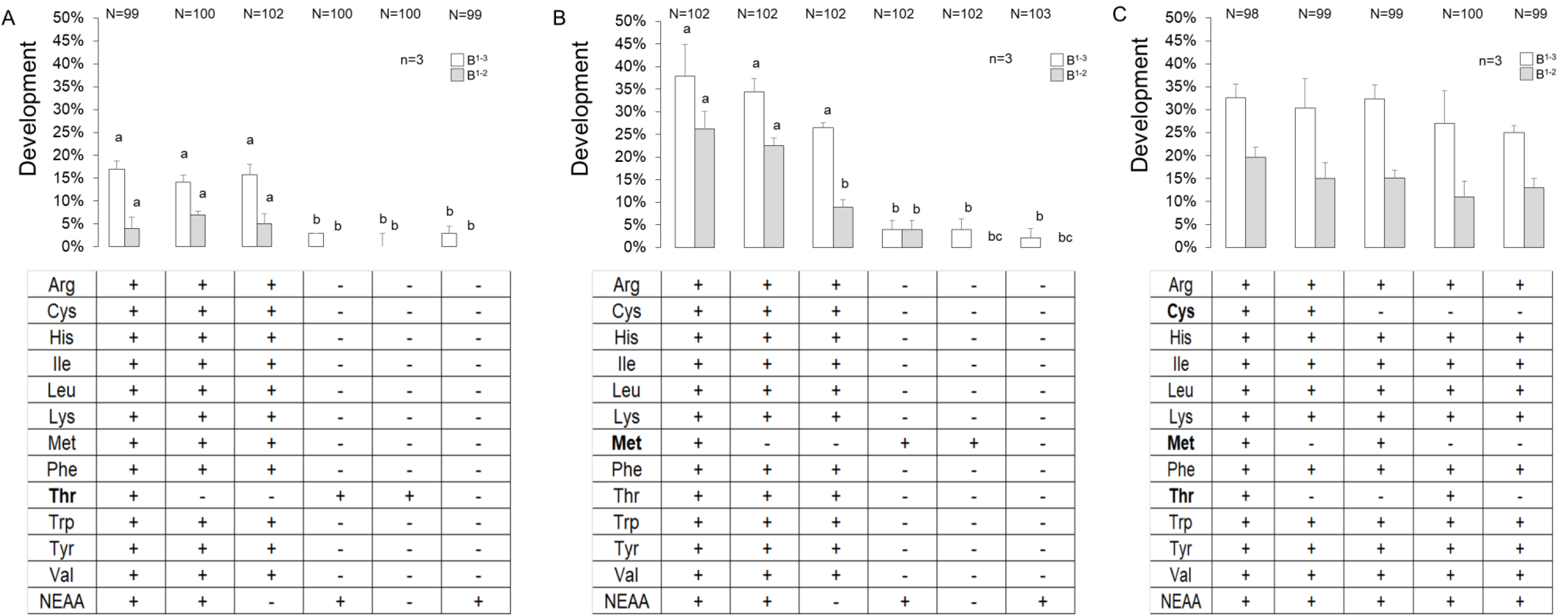
Effect of candidate amino acid dropout on bovine blastocyst development. Effect of depleting (A) threonine (-T), (B) methionine (-M) or (C) combinations of threonine, methionine and cysteine (-MT, -CM, -CT, -CMT) on blastocyst development into B1-3 and B1-2. EAA and NEAA depletion, with or without providing threonine or methionine (+T, +M) served as additional treatments. ab= groups with different letters within each grading category differ P<0.005. Error bars = sem; n = no. of independent IVF experiments (N = no. of embryos placed into IVC); Dropped out amino acids (bold) are shown in 3 letter code; NEAAs = non-essential AAs.

Human PSCs do not survive without methionine [19]. By contrast, bovine blastocyst development was not affected when embryos were deprived of methionine (Fig. 1B). Again, methionine provision did not rescue blastocyst development in the absence of other EAAs. Even combined threonine, methionine and cystine dropout (‘-MT’, ‘-CT’, ‘-CM’, ‘-CMT’) had no significant effect (Fig. 1C), indicating that key EAAs involved in the threonine-SAM pathway are dispensable for the morula-to-blastocyst transition.

Since lysine and leucine are also critical for human PSC development [19], we tested their effect using the same experimental regime as before. Together with closely related isoleucine, we performed dual (‘-LK’, ‘-IL’, ‘-IK’) and triple (‘-ILK’) candidate EAA dropout experiments. Blastocyst development was not affected in any of these combinations (Fig. S1D). By contrast, dropout of all six of these EAA candidates (‘-CILKMT’) significantly reduced development (Fig. S1E). This reduction was stronger than dropping out the remaining other six EAAs (’-FHRYVW). Removing each group of six EAAs still resulted in better development than removing all 12 EAAs (Fig. S1E). No rescue was observed when nine EAAs were dropped out and either cystine, methionine and threonine (‘+CMT’) or isoleucine, leucine and lysine (‘+ILK’) were provided (Fig. S1F). In summary, neither threonine nor methionine, leucine or lysine dependency could be confirmed in cultured bovine embryos. As no other clear candidates emerged from this dropout screen, we decided to directly test the role of threonine catabolism.

### TDH is expressed in bovine blastocysts

Mouse ICM cells and ESCs, their *in vitro* counterpart, abundantly express the enzyme TDH which converts threonine into glycine and acetyl-CoA. We determined *TDH* expression in various bovine embryonic and adult tissues by qRT-PCR (Fig. 2A). The gene was detected in blastocysts where it showed no differential expression between ICM and TE cells. Its expression levels were lower in liver and testis tissue samples and significantly reduced in primary BEFs. None of various differentiated adult tissues (brain, heart, kidney, muscle, spleen and skin) showed detectable *TDH* expression, similarly to what has been described in mouse [13].

**Figure 2.**
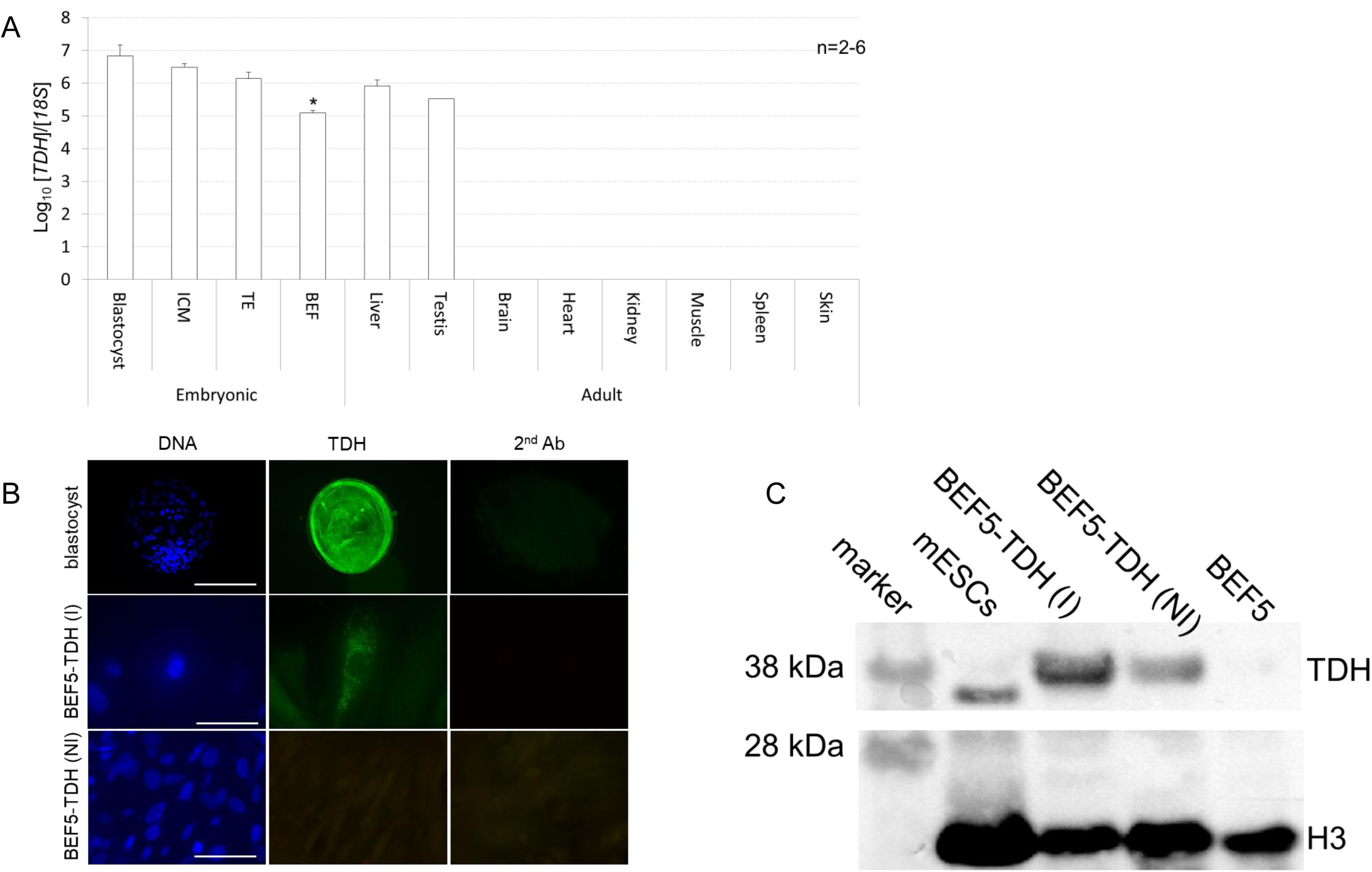
TDH expression in bovine cells. A) qRT-PCR for *TDH* gene expression in bovine embryonic and adult tissues. Relative values were normalised on *18S* expression. Error bars = sem. * = Groups differ P<0.05 from the ICM reference group; n = no. of independent biological replicates. B) Immunofluorescence staining of TDH in bovine embryonic cells. Images were pseudo-colored using ImageJ. DNA was stained with Hoechst 33342 (blue), TDH was stained with Alexa Fluor® 488 (green). In the “2^nd^Ab” panel, primary antibody incubation was omitted. Scale bar = 100 μm. C) Immunoblot of TDH-overexpressing bovine cells. Ectopic TDH_MYC was detected with an anti-TDH antibody in TDH-transgenic cells (BEF-TDH) but not in the parental line only carrying the Tet3G reverse tetracycline transactivator (BEF5). Mouse embryonic stem cells (mESCs) provide a positive control. Histone 3 (H3) was used as a loading control. The exposure time was 30 seconds. I = Induced, NI = Non-induced

In eukaryotic cells, the TDH enzyme is located within mitochondria where it supplies acetyl-CoA for the Krebs cycle and glycine to tetrahydrofolate charging [29]. We first confirmed by qRT-PCR that *Tdh* mRNA is copiously expressed in mouse ESCs and blastocysts but not detectable in mouse embryonic fibroblasts (Fig. S2A). Next we determined localisation and abundance of the TDH protein by immunofluorescence and immunoblotting, respectively. Using polyclonal antibodies against the mouse protein [13], we detected punctate staining in the cytoplasm of ESCs, resembling the perinuclear distribution of mitochondria (Fig. S2B). TDH protein was also detected by western blotting in murine iPSCs but absent in MEFs and adult muscle fibroblasts (Fig. S2C).

The polyclonal antisera used for these experiments were raised against two different synthetic peptides derived from the mouse protein (Fig. S3A). The peptides were 90% (2/21 mismatches) and 76% (5/21 mismatches) identical with the bovine TDH sequence. To validate specific antibody cross-reactivity with bovine, a transgenic *TDH*-overexpressing bovine embryonic fibroblast line (BEF5-TDH) was derived. This line expressed the full-length TDH protein, carrying a C-terminal MYC-tag, under control of a Dox-inducible tetracycline response element (Fig. S3B). Immunostaining against TDH_C-MYC with a C-MYC tag antibody showed strong signals only in Dox-induced BEF5-TDH cells but not in non-induced controls, co-localising with TDH antibody signals (Fig. S3C). This predominantly perinuclear immunoreactivity co-localised with mitotracker signals, indicating that the expected subcellular localisation of TDH was not altered by the presence of a MYC-tag (Fig. S3D).

As expected from the mRNA expression, TDH protein was homogeneously detectable in the ICM and TE of bovine blastocysts (Fig. 2B), as well as in mouse blastocysts stained as a positive control (data not shown). Specific detection of bovine TDH was confirmed by western blotting with the polyclonal antisera (Fig. 2C). A band around the expected molecular weight (41 kDa) was detected in Dox-induced TDH-overexpressing BEF5 cells but not in the parental line only carrying the Tet3G reverse tetracycline transactivator (BEF5). Non-induced BEF5 showed reduced expression levels, possibly due to leaky expression from the TRE3G promoter, while mouse ESCs showed a slightly lower band of endogenous TDH without the MYC-tag.

### TDH inhibition alters gene expression and compromises blastocyst development

We next examined the effect of pharmacological TDH inhibition in functional studies. We employed Qc1, a reversible, non-competitive TDH inhibitor that blocks the enzyme from catabolising threonine into glycine and acetyl-CoA. First, we compared the effect of different Qc1 concentrations (1, 5, 10, 50, 67, 100 µM) against their respective solvent controls on blastocyst development. From this, the minimal effective dose was determined as 50 µM (data not shown). At this concentration Qc1 completely abolished mouse blastocyst development (Fig. S4A, B). In bovine, treatment from D5 to D8 significantly reduced total development from 37% to 26% and more than halved the output of high-quality blastocysts from 18% to 8% (*P*<0.005, Table 1). This was accompanied by altered gene expression of ICM-and TE-specific marker genes. Specifically, ICM-restricted genes (*NANOG, FGF4, SOX2*) tended to be up-regulated, while TE-restricted genes were down-regulated (*CDX2, KRT8*) compared to DMSO solvent controls (Fig. 3).

**Figure 3.**
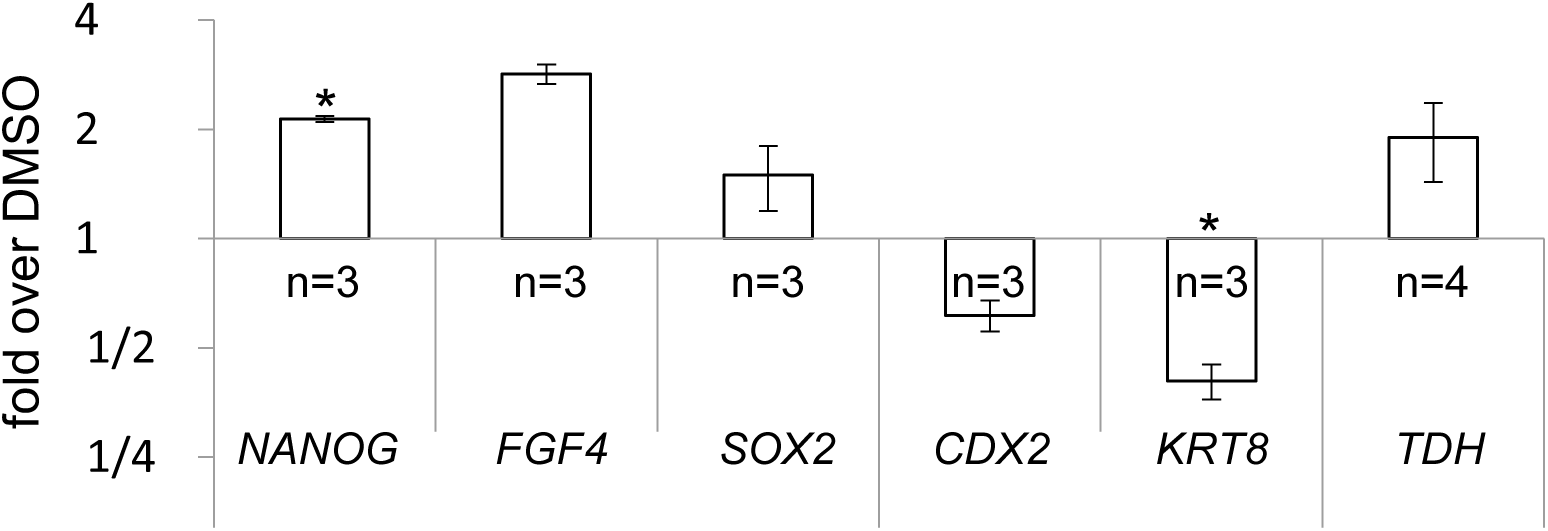
Gene expression changes induced by TDH inhibition. qRT-PCR for ICM (*NANOG, FGF4, SOX2*) versus TE (*CDX2, KRT8*) marker genes and *TDH*. cDNA was extracted from pools of grade 1-2 D8 blastocysts cultured in DMSO-vs. Qc1-containing medium. Target gene values were normalized on 18S expression and expressed as fold-change over DMSO control; * bars differ P<0.05 from DMSO control; Error bars = sem. n = no. of independent biological replicates.

**Figure 4.**
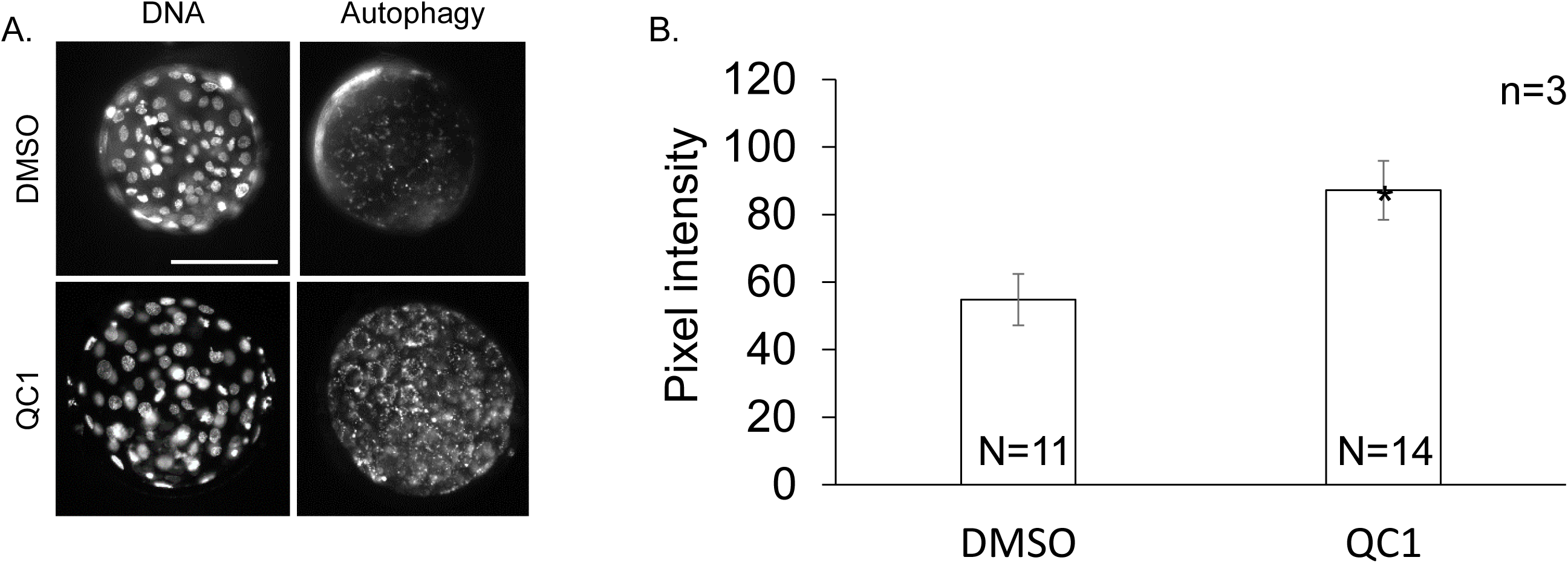
TDH inhibition increases autophagy in bovine embryos. A) IVF-produced embryos were treated with DMSO (vehicle control) versus TDH inhibitor Qc1 from the morula-stage (D5) onwards. Fluorescence signal from autophagic vacuoles was detected in D8 blastocysts. B) Fluorescence intensity on representative blastocyst images was quantified with ImageJ. n= no. of biological replicates. Error bars = sem; * = group differs P<0.05 from DMSO control by t-test; N= no. of quantified embryos.

Expression of *TDH* itself was not significantly altered by its pharmacological inhibition. Despite these changes in lineage-specific marker gene expression, the ratio of inner (ICM) to outer (TE) cells, as determined by differential staining (Fig. S5) was not affected by TDH inhibition (Table 2). However, there was an almost 2-fold decrease in ICM, TE and total cell numbers (1.9-, 1.8-and 1.8-fold, respectively, *P*<0.005), in line with the observed 2.25-fold decrease of morphologically high-grade blastocysts.

**TABLE 2.**
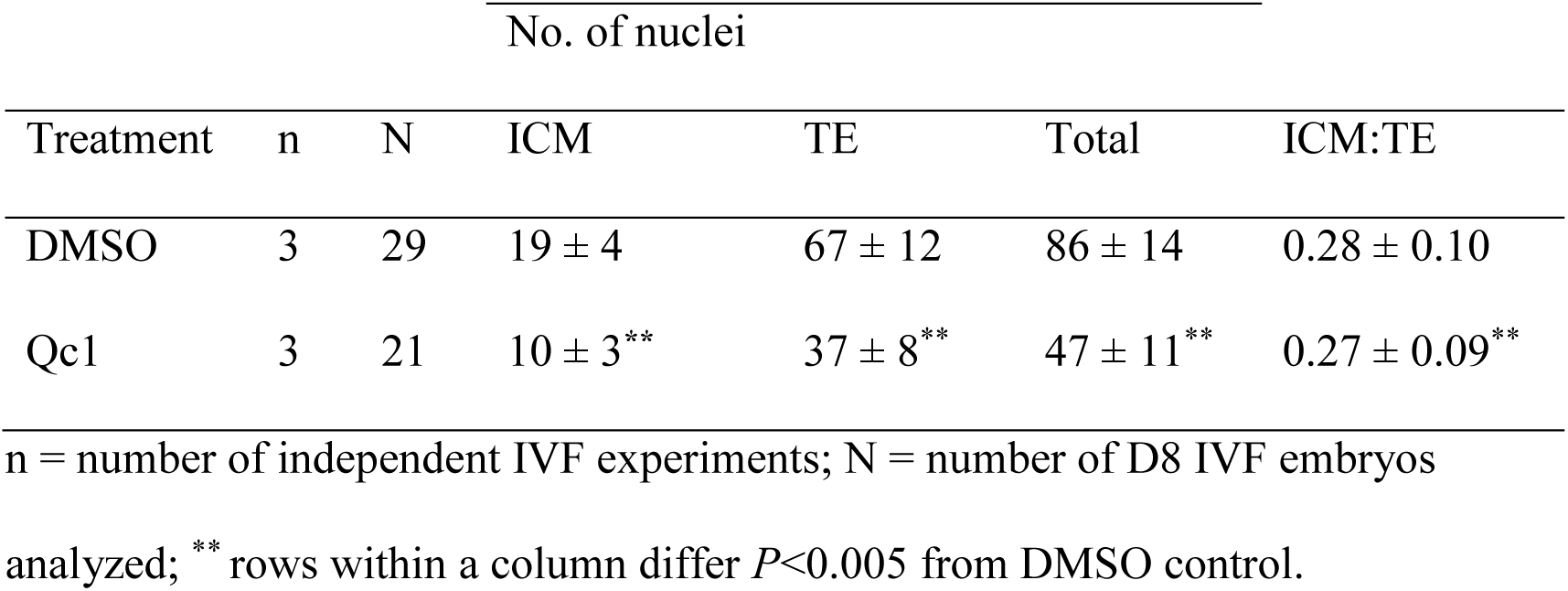
Effect of TDH inhibition on ICM and TE nuclei numbers (mean ± SD)

### TDH inhibition increases autophagy in bovine embryos

The reduced cell number in Qc1-treated embryos could be due to increased cell death as TDH has been described to specifically stimulate autophagy in ESCs [16]. Using a commercial detection kit, autophagy was detected and quantified in D8 bovine embryos. In the Qc1-treated group, the fluorescent signal associated with autophagic vacuoles (Fig. 3A) increased 1.5 fold compared to DMSO controls (P<0.05), providing a potential mechanism for the reduction in blastocyst cell numbers upon TDH inhibition (Fig. 3B).

## DISCUSSION

Here we show that threonine catabolism was required for bovine blastocyst viability. In its absence, embryo survival was compromised due to increased autophagy in both TE and ICM. This observation complements findings in mouse, where threonine catabolism appeared to be particularly critical for development of the ICM and ICM-derived PSCs.

### Amino acid dependency during bovine embryo development

We undertook a targeted amino acid dropout screen on cattle embryos to investigate blastocyst development in the absence of individual amino acids. For this, we first removed potentially contaminating amino acid sources by depleting the base medium of BSA. On its own, BSA does not appear to alter the concentration of amino acids in the medium [30]. However, BSA is taken up by embryos through fluid-phase endocytosis [10, 31]. Uptake and subsequent intracellular degradation probably maintain intracellular amino acid pools [10]. Therefore, BSA is an undefined amino acid source that could confound experiments examining the role of amino acid metabolism *in vitro*. In pilot experiments, we established that BSA or amino acid dropout from D7 to D8 had no effect on development (data not shown). By contrast, BSA depletion from D5 to D8 modestly compromised blastocyst development, even in the presence of glutamine, NEAAs and EAAs, supporting a nutritive role of BSA [32]. Although albumin is not an absolute requirement for bovine embryo development *in vitro* [33], its inclusion in the culture medium has been generally considered beneficial [32]. This benefit may be particularly important post-compaction when the embryo increases total protein content via increased protein uptake [31], coinciding with net cell growth and proliferation.

To further refine a minimal medium for dropout experiments, we also omitted glutamine from the BSA-free medium. There is conflicting information regarding the requirement of this amino acid from the morula stage onwards. Whilst it may not be required for blastocyst quality, glutamine depletion from BSA-, NEAA-and EAA-containing medium reduced blastocyst development [34]. This supports the finding that increasing glutamine uptake in expanding and hatching blastocysts feeds as an energy substrate into the Krebs cycle [35-38]. However, others have not found significant glutamine depletion from the medium by bovine [8] or mouse [39] blastocysts. We did not directly address the role of glutamine in our study.

It was further necessary to wash D5 embryos in protein-free buffer before transferring them into the amino acid dropout medium. Without this step, removing all NEAAs or EAAs on D5 had no significant effect on development in the absence of BSA. By contrast, washing embryos before transfer into dropout medium unmasked the detrimental effect of AA removal. We reasoned that this may have been due to residual protein carry-over from embryo-conditioned BSA-containing ESOF to fresh BSA-free LSOF.

As a last step to exclude contaminating amino acid sources, we cultured embryos individually to better detect potential amino acid dropout effects. Under group culture conditions, removing all NEAAs and/or EAAs on D5 had no significant effect on blastocyst development in the absence of BSA. Once embryos were cultured individually without protein carry-over, dropping out BSA strongly reduced blastocyst development, indicating that the embryos themselves can be a potentially confounding source of amino acids.

We observed that individually cultured bovine embryos without BSA, glutamine and protein carry-over required both NEAAs and EAAs for maximal blastocyst development. This confirmed earlier findings, obtained in BSA-containing medium, that the combination of all 20 amino acids stimulated bovine blastocyst development and quality [34]. This earlier study also found that NEAAs supported the highest proportion of bovine ICM cells, in contrast to mouse, where EAAs stimulated ICM development [40]. We did not quantify cell numbers but observed that the absence of NEAAs vs EAAs, reduced total blastocyst development by one-third vs two-third, respectively. Since lack of EAAs had the most detrimental effect on blastocyst development, we decided to deprive individually cultured embryos of candidate EAAs from D5 to D8 in subsequent experiments.

### Threonine dependency during bovine embryo development

Using systematic depletion of all 20 amino acids from cell culture medium, threonine was identified as the only critical amino acid in for growing mouse ESCs [13]. Without threonine, thymidine biosynthesis and ESC self-renewal was severely impaired. Mouse blastocysts do not deplete large amounts of threonine from the medium [39] and, to our knowledge, a detrimental effect of threonine dropout on mouse blastocyst development has not yet been conclusively demonstrated. Here we used a similar experimental dropout approach to show that total development of bovine blastocysts was not affected by the absence of threonine. This is despite the finding that bovine embryos consume significant amounts of threonine at all stages of development *in vitro*, in particular at the blastocyst stage [8]. In fact, threonine was the only amino acid that showed this particular depletion pattern, consistent with our finding that threonine dropout compromised blastocyst development. In particular, isolated ICM depleted threonine (together with asparagine, glycine, tyrosine, tryptophan and phenylalanine), even though this effect was not obvious in intact blastocysts [41]. Cultured human PSCs do not survive without lysine, leucine or methionine [19]. In cultured bovine embryos, on the other hand, methionine dropout did not affect development, consistent with its lack of net uptake in isolated ICMs and intact blastocysts at this stage [41]. Blastocyst development was also not affected when embryos were simultaneously deprived of two or three of these EAAs. Thus key EAAs involved in mouse and human embryonic PSC survival are dispensable for bovine blastocyst and ICM development under the tested experimental conditions. As no promising candidate EAA emerged from this dropout screen, we directly tested threonine dependency with a different experimental approach.

### Role of TDH in regulating pluripotency-related molecular pathways

Mouse TDH mRNA and protein expression, respectively, was restricted to the ICM [13]. By contrast, isolated bovine ICM and TE showed no significant enrichment of *TDH* mRNA in either tissue. Using Dox-inducible overexpression of MYC-tagged bovine TDH, we found that polyclonal anti-mouse TDH antisera specifically recognised a protein of the correct molecular weight (∼42 kDa) in the mitochondria of BEFs. However, immunofluorescence with this validated TDH antibody showed strong homogeneous immunoreactivity across the whole bovine blastocyst. Thus TDH expression was not ICM-restricted in cattle. This is in contrast to the general conservation of ICM-restricted key pluripotency markers between mouse and bovine, including phosphorylated STAT3, SOX2 and NANOG [28]. Expression of other pluripotency-related transcription factors, such as *POU5F1* and *KLF4*, also extends into the bovine TE [42], suggesting that some aspects of pluripotency acquisition may be delayed in cattle compared to mouse.

In mouse, metabolic remodelling plays an important function in regulating somatic cell reprogramming into pluripotency. This is supported by solid evidence that the transition from oxidative phosphorylation to glycolysis [43, 44], as well as TDH-mediated threonine catabolism promote iPSC formation. The detailed mechanism of this process remains to be elucidated but appears to involve post-transcriptional and post-translational regulation by microRNA miR-9 and protein arginine methyltransferase PRMT5, respectively [17]. Specifically, miR-9 represses TDH expression by targeting its 3’UTR and inhibiting TDH-facilitated reprogramming. PRMT5, a direct TDH binding partner, increases TDH enzyme activity and TDH-enhanced reprogramming efficiency [45]. Furthermore, pluripotency genes (*Pou5f1, Sox2, Nanog, Zfp42 and Blimp1*) were down-regulated upon TDH depletion in mouse ESCs [15]. This role of TDH as a positive regulator of pluripotency was seemingly at odds with the modest up-regulation of pluripotency-associated genes (*NANOG, FGF4, SOX2*) upon TDH inhibition. It remains to be seen if the interactions described for iPS and ESC reprogramming *in vitro* also play a physiological role in establishing pluripotency during bovine embryogenesis.

DNA and histone methylation is a major regulator of gene expression. Both modifications are catalysed by methyltransferases that use SAM as a methyl group donor and SAM levels were elevated in mouse PSCs compared to fibroblasts [15]. This increased accumulation of SAM was generated by uptake of extracellular threonine and Tdh activity to fuel histone methylation, specifically of histone 3 lysine 4 di-and trimethylation (H3K4me2, H3K4me3). In human PSCs, methionine specifically contributed to SAM production, resulting in increased H3K4me3, while other trimethylation markers (H3K9me3 and H3K27me3) were not affected [12]. Thus threonine-dependent synthesis of SAM and high SAM/SAH ratios correlated with H3K4me2/3 levels, revealing a possible mechanistic link between metabolic and epigenetic states. It remains to be determined if threonine restriction or Qc1 treatment could elicit a similarly specific response of SAM/SAH ratios and H3 modifications in bovine embryos.

Another pluripotency pathway regulating SAM-dependent histone methylation is leukaemia inhibitory factor (LIF)/STAT signalling. In human PSCs, LIF/STAT signals upregulated nicotinamide N-methyltransferase (*NNMT*), a major consumer of SAM [50]. The ensuing reduction in SAM levels correlated with decreased levels of repressive H3K9me3 and H3K27me3. This, in turn, repressed hypoxia-inducible factor (HIF) 1α and promoted canonical WNT signalling, delaying exit from naive pluripotency. As STAT3-signals have been shown to be restricted to the bovine ICM [28], it would be informative to investigate if such an STAT-NNMT-SAM-H3K27me3-HIF/WNT-axis was also operational in bovine blastocysts. This could help to uncover potentially species-specific connections between metabolism, epigenetics and embryonic development.

### TDH regulating autophagy and embryonic cell survival

To determine if TDH was not only present in bovine embryos but also functional during development, we used pharmacological inhibition with Qc1 [16]. This compound reversibly blocks the ability of the TDH enzyme to catabolize threonine into glycine and acetyl-CoA, cutting off the supply for nucleotide synthesis and ATP generation, respectively [29]. In ESCs, chemical inhibition of TDH enzyme activity prompted cellular autophagy, an intracellular bulk degradation process that is activated by various stressors, including restriction of nutrient supply. The same chemical inhibitor displayed no toxicity to somatic cells that do not express the TDH enzyme. We first confirmed that Qc1 was specifically killing embryonic cells. Both ESCs (data not shown) and mouse embryos did not survive continuous exposure to 50 μM QC1. This confirmed and extended earlier findings that Qc1 is toxic to embryonic mouse cells. For treatment of bovine cells, we established a dose-response curve by treating D5 IVF embryos with 1, 5, 10, 50, 66.7 and 100 μM Qc1. Compared to equivalent DMSO solvent controls, 1-10 μM had no detectable effect while 50-100 μM significantly reduced total and transferable grade D8 embryo development (data not shown). At the effective concentration of 50 μM, proliferation and survival of BEF cells was not obviously compromised, indicating that cells with lower TDH expression levels may be less susceptible to the detrimental effects of Qc1. Overall blastocyst development was reduced and the fewer blastocysts that formed were of disproportionately poorer quality, resulting in less than half the number of transferable grade embryos. These morphological observations correlated well with differential stain quantification, showing an almost 2-fold decrease in both ICM and TE cell numbers. This equal reduction in cell numbers from both lineages also correlated well with overall increased levels of autophagy in blastocysts. Distribution of the fluorescent signal from autophagic vacuoles did not show any obvious differences between ICM and TE cells, indicating that autophagy was equally increased in both cell populations. This observation was again consistent with the similar TDH expression levels, both for mRNA and protein, in ICM and TE cells.

Autophagy is a conserved catabolic process that recycles cytoplasmic material during low energy conditions. It naturally occurs during early preimplantation development in bovine, consistent with the high relative abundance of autophagy-associated transcripts in oocytes and early embryos up to the 4-cell stage, but low levels at the morula-to-blastocyst stages [51]. In mouse embryos, autophagy also declined rapidly after the 4-cell stage [52]. This early autophagy appeared to be beneficial and stimulating it pharmacologically improved development and quality of IVP bovine blastocysts. However, the time point at which stimulation of autophagic activity occurs was critical. If autophagy was stimulated late, i.e., during the last 4 days of *in vitro* development, it reduced blastocyst development rate, as well as ICM and TE cell numbers [51]. Consistent with our results, these findings indicate that high autophagic activity is essential during early pre-implantation development but detrimental during later stages. This would have to be confirmed by additional indicators of hyperautophagy in Qc1-treated embryos, such as increased LC3-I to LC3-II conversion, dot-like LC3 immunoreactivity and direct ultrastructural detection of autophagosomes. Cellular autophagy precedes loss of viability in other cell types, such as mouse ESCs [16]. In bovine embryos, the Qc1-induced autophagic state was also insufficient to prevent cell death. These results confirm the importance of the threonine-dependent metabolic state for bovine blastocyst viability. They also highlight that metabolic specialization of a particular cell type, such as mouse ICM cells and ICM-derived PSCs, is not necessarily conserved between species. Future experiments will need to better characterise the metabolome of bovine ICM vs TE cells in order to define a unique metabolic state that may aid in the derivation of bovine embryonic PSCs.

## ACKNOWLEDGEMENTS

We are grateful to Dr Jingwei Wei for excellent technical help with mouse embryology. We thank Drs Rudolf Jaenisch and Steven McKnight for donating v6.5 ESCs and TDH antibodies, respectively. We acknowledge the Wellcome Trust Sanger Institute for making *pCyL43* transposase available. This work was funded by MSI contracts C10X0303, C10X1002 and AgResearch.

